# A systems-level analysis highlights microglial activation as a modifying factor in common forms of human epilepsy

**DOI:** 10.1101/470518

**Authors:** Andre Altmann, Mina Ryten, Martina Di Nunzio, Teresa Ravizza, Daniele Tolomeo, Regina H Reynolds, Alyma Somani, Marco Bacigaluppi, Valentina Iori, Edoardo Micotti, Juan A. Botía, Julie Absil, Saud Alhusaini, Marina K. M. Alvim, Pia Auvinen, Nuria Bargallo, Emanuele Bartolini, Benjamin Bender, Felipe P. G. Bergo, Tauana Bernardes, Andrea Bernasconi, Neda Bernasconi, Boris C. Bernhardt, Karen Blackmon, Barbara Braga, Maria Eugenia Caligiuri, Anna Calvo, Chad Carlson, Sarah J. Carr, Gianpiero L. Cavalleri, Fernando Cendes, Jian Chen, Shuai Chen, Andrea Cherubini, Luis Concha, Philippe David, Norman Delanty, Chantal Depondt, Orrin Devinsky, Colin P. Doherty, Martin Domin, Niels K. Focke, Sonya Foley, Wendy Franca, Antonio Gambardella, Renzo Guerrini, Khalid Hamandi, Derrek P. Hibar, Dmitry Isaev, Graeme D. Jackson, Neda Jahanshad, Reetta Kalviainen, Simon S. Keller, Peter Kochunov, Raviteja Kotikalapudi, Magdalena A. Kowalczyk, Ruben Kuzniecky, Patrick Kwan, Angelo Labate, Soenke Langner, Matteo Lenge, Min Liu, Pascal Martin, Mario Mascalchi, Stefano Meletti, Marcia E. Morita, Terence J. O’Brien, Jose C. Pariente, Mark P. Richardson, Raul Rodriguez-Cruces, Christian Rummel, Taavi Saavalainen, Mira K. Semmelroch, Mariasavina Severino, Pasquale Striano, Thomas Thesen, Rhys H. Thomas, Manuela Tondelli, Domenico Tortora, Anna Elisabetta Vaudano, Lucy Vivash, Felix von Podewils, Jan Wagner, Bernd Weber, Roland Wiest, Clarissa L. Yasuda, Guohao Zhang, Junsong Zhang, ENIGMA-Epilepsy Working Group, Costin Leu, Andreja Avbersek, EpiPGX Consortium, Maria Thorn, Christopher D Whelan, Paul Thompson, Carrie McDonald, Annamaria Vezzani, Sanjay M Sisodiya

**Affiliations:** Centre for Medical Image Computing, University College London, London, UK; Department of Neurodegenerative Disease, UCL Queen Square Institute of Neurology, London, UK; Department of Neuroscience, Istituto di Ricerche Farmacologiche Mario Negri IRCCS, Milano, Italy; Division of Neuropathology, UCL Queen Square Institute of Neurology, London, UK; Department of Neurology, San Raffaele Scientific Institute and Vita Salute San Raffaele University, Milan, Italy; Departamento de Ingeniería de la Informatión y las Comunicaciones. Universidad de Murcia, Murcia, Spain; Department of Radiology, Hôpital Erasme, Universite Libre de Bruxelles, Brussels 1070, Belgium; Department of Molecular and Cellular Therapeutics, Royal College of Surgeons in Ireland, Dublin, Ireland; Department of Neurology and Neurosurgery, Montreal Neurological Institute, McGill University, Montreal, Quebec, Canada; Department of Neurology, University of Campinas, Campinas, Brazil; Epilepsy Center, Department of Neurology, Kuopio University, Kuopio, Finland; Institute of Clinical Medicine, Neurology, University of Eastern Finland, Kuopio, Finland; Magnetic Resonance Image Core Facility, IDIBAPS, Barcelona, Spain; Centre de Diagnostic Per la Imatge (CDIC), Hospital Clinic, Barcelona, Spain; Pediatric Neurology Unit, Children’s Hospital A. Meyer-University of Florence, Italy; IRCCS Stella Maris Foundation, Pisa, Italy; Department of Diagnostic and Interventional Neuroradiology, University of Tübingen, Tübingen, Germany; Neuroimaging of Epilepsy Laboratory, Montreal Neurological Institute and Hospital, McGill University, Montreal, Quebec, Canada; Multimodal Imaging and Connectome Analysis Lab, Montreal Neurological Institute and Hospital, McGill University, Montreal, Quebec, Canada; Comprehensive Epilepsy Center, Department of Neurology, New York University School of Medicine, New York, USA; Department of Physiology, Neuroscience and Behavioral Science, St. George’s University, Grenada, West Indies; Institute of Molecular Bioimaging and Physiology of the National Research Council (IBFM-CNR), Catanzaro, Italy; Medical College of Wisconsin, Department of Neurology, Milwaukee, Wl, USA; Department of Basic and Clinical Neuroscience, Institute of Psychiatry, Psychology and Neuroscience, King’s College London, UK; FutureNeuro Research Centre, RCSI, Dublin, Ireland; Department of Computer Science and Engineering, The Ohio State University, USA; Cognitive Science Department, Xiamen University, Xiamen, China; Fujian Key Laboratory of the Brain-like Intelligent Systems, China; Instituto de Neurobiología, Universidad Nacional Autónoma de México. Querétaro, Querétaro, México; Division of Neurology, Beaumont Hospital, Dublin 9, Ireland; Department of Neurology, Hôpital Erasme, Universite Libre de Bruxelles, Brussels 1070, Belgium; Neurology Department, St. James’s Hospital, Dublin 8, Ireland; Functional Imaging Unit, Institute of Diagnostic Radiology and Neuroradiology, University Medicine Greifswald, Greifswald, Germany; Department of Neurology and Epileptology, Hertie Institute for Clinical Brain Research, University of Tübingen, Tübingen, Germany; Department of Clinical Neurophysiology, University Medicine Göttingen, Göttingen, Germany; Cardiff University Brain Research Imaging Centre, School of Psychology, Wales, UK; Institute of Neurology, University, “Magna Grascia”, Catanzaro, Italy; Institute of Psychological Medicine and Clinical Neurosciences, Hadyn Ellis Building, Maindy Road, Cardiff, UK; Department of Neurology, University Hospital of Wales, Cardiff, UK; Imaging Genetics Center, Mark and Mary Stevens Neuroimaging and Informatics Institute, University of Southern California, Los Angeles, California, USA; The Florey Institute of Neuroscience and Mental Health, Austin Campus, Melbourne, VIC, Australia; Florey Department of Neuroscience and Mental Health, The University of Melbourne, Melbourne, VIC, Australia; Department of Molecular and Clinical Pharmacology, Institute of Translational Medicine, University of Liverpool, UK; Maryland Psychiatric Research Center, Department of Psychiatry, University of Maryland School of Medicine, Maryland, USA; Department of Neurology, Royal Melbourne Hospital, Parkville, 3050, Australia; Neuroradiology Unit, Children’s Hospital A. Meyer, Florence, Italy; “Mario Serio” Department of Experimental and Clinical Biomedical Sciences, University of Florence, Italy; Department of Biomedical, Metabolic, and Neural Science, University of Modena and Reggio Emilia, NOCSE Hospital, Modena, Italy; Department of Medicine, University of Melbourne, Parkville, VIC, 3052, Australia; Department of Neurology, King’s College Hospital, London, UK; Support Center for Advanced Neuroimaging (SCAN), University Institute for Diagnostic and Interventional Neuroradiology, Inselspital, University of Bern, Bern, Switzerland; Central Finland Central Hospital, Medical Imaging Unit, Jyväskylä, Finland; Neuroradiology Unit, Department of Head and Neck and Neurosciences, Istituto Giannina Gaslini, Genova, Italy; Pediatric Neurology and Muscular Diseases Unit, Department of Neurosciences, Rehabilitation, Ophthalmology, Genetics, Maternal and Child Health, University of Genoa, Genova, Italy; Melbourne Brain Centre, Department of Medicine, University of Melbourne, Parkville, VIC, 3052, Australia; Department of Neurology, University Medicine Greifswald, Greifswald, Germany; Department of Epileptology, University Hospital Bonn, Bonn, Germany; Department of Neurology, Philips University of Marburg, Marburg Germany; Department of Neurocognition / Imaging, Life & Brain Research Centre, Bonn, Germany; Department of Computer Science and Electrical Engineering, University of Maryland, Baltimore County, USA; Genomic Medicine Institute, Lerner Research Institute, Cleveland Clinic, Cleveland, OH, USA; Stanley Center for Psychiatric Research, Broad Institute of Harvard and MIT, Cambridge, MA, USA; Department of Clinical and Experimental Epilepsy, UCL Institute of Neurology, London, UK; Multimodal Imaging Laboratory, University of California San Diego, San Diego, California, USA; Department of Psychiatry, University of California San Diego, San Diego, California, USA; Chalfont Centre for Epilepsy, Bucks, UK

## Abstract

The common human epilepsies are associated with distinct patterns of reduced cortical thickness, detectable on neuroimaging, with important clinical consequences. To explore underlying mechanisms, we layered MRI-based cortical structural maps from a large-scale epilepsy neuroimaging study onto highly spatially-resolved human brain gene expression data, identifying >2,500 genes overexpressed in regions of reduced cortical thickness, compared to relatively-protected regions. The resulting set of differentially-expressed genes shows enrichment for microglial markers, and in particular, activated microglial states. Parallel analyses of cell-specific eQTLs show enrichment in human genetic signatures of epilepsy severity, but not epilepsy causation. *Post mortem* brain tissue from humans with epilepsy shows excess activated microglia. In an experimental model, depletion of activated microglia prevents cortical thinning, but not the development of chronic seizures. These convergent data strongly implicate activated microglia in cortical thinning, representing a new dimension for concern and disease modification in the epilepsies, potentially distinct from seizure control.

## Introduction

Significant progress is being made in understanding disease processes in the epilepsies. Many genetic variants causing or associated with rare and common epilepsies have been reported^1,2^, with discovery continuing apace^3^. Numerous direct structural causes of epilepsy have been revealed by brain magnetic resonance imaging (MRI), since its widespread implementation in the period immediately preceding the extensive application of genetics. Several structural abnormalities are now themselves known to have a genetic basis^4^. As a result, the proportion of causally-explicable epilepsies is growing rapidly. Conversely, the mechanisms whereby these identified causes promote epileptogenesis and seizures remain obscure for most human epilepsies. Moreover, beyond causation and epileptogenesis, the epilepsies involve many other processes: some lead to clinically-apparent consequences, such as developmental delay or memory dysfunction, whereas others, without necessarily obvious symptoms, may be detected only on investigation - cerebellar atrophy is one example. The natural history of any given epilepsy need not be a single linear dynamic from causation to a unique, predictable, final outcome: for example, the epilepsies are associated with shortened longevity (even if seizures stop)^5^ and increased risk of particular comorbidities^6^. Known causes *per se* may not explain all the observed outcomes, suggesting that many epilepsies could be conceptualised as intricate matrices of causation, processes and outcomes^7^, with complex inter-dependencies, such as a likely link between reduction in cortical thickness and disease duration^8^.

Through the ENIGMA-Epilepsy consortium, we recently showed that across a wide range of common human epilepsies, which are known to have both distinct and shared genetic architecture^2,3,9,10^, there are also shared, pan-syndrome, and distinct, syndrome-specific, regional patterns of altered cortical thickness and altered subcortical grey matter volumes^8^. The causes of the structural changes in these epilepsies are not known. The findings suggest structural losses may reflect an initial insult, subsequent epileptogenesis or progressive neurodegeneration, or some combination, and indicate that the epilepsies cannot necessarily be considered entirely benign at the structural level. We sought to identify processes underlying the structural findings.

Human neurological disease biology has been successfully revealed using powerful combinations of brain MRI findings, regionalised brain-specific gene expression and gene co-expression network datasets, in systems biology approaches implicating candidate genes^11–19^. Here, we used the findings from the ENIGMA-Epilepsy study to direct interrogation for regionalised gene expression changes, which in turn identified candidate genes which might underlie thickness or volume reductions across the studied epilepsy syndromes. Gene expression is itself largely driven by underlying constitutional genetic variation. We therefore hypothesised that this approach could pick out disease mechanisms causing the observed structural changes. We then tested whether the postulated mechanisms were identifiable from epilepsy susceptibility or treatment-response genome-wide association studies and further explored the findings with a series of additional experiments in both human and animal tissues. We demonstrate experimentally that depletion of the cell type, microglia, most strongly implicated by the systems biology approach can successfully avert cortical thinning. Together, the results point to the potential for prevention of the widespread, but largely unstudied, reduction in cortical thickness that accompanies common human epilepsies.

## Results

### Cortical regions at most risk of reduced thickness in individuals with common forms of epilepsy are characterised by higher expression of over 2500 genes

Through the ENIGMA-Epilepsy consortium, we recently showed that across a wide range of common epilepsies, there are distinctive regional patterns of altered cortical thickness and altered subcortical grey matter volumes^8^. To identify the molecular basis of this regional vulnerability in the broad spectrum of epilepsies, we focused our analyses on the shared pan-syndrome cortical findings and leveraged data from the Allen Institute for Brain Science (AIBS) healthy human brain gene expression atlas (Figure 1). We chose to study all the epilepsies as a group not only because of the general importance of pan-syndrome cortical findings, but because these structural changes were most likely to reveal the condition-specific, as opposed to syndrome-specific, processes driving the selective vulnerability of cortical regions. We hypothesised that matching patterns of regional cortical vulnerability to reduced thickness with patterns of gene expression differences across the human cortex would point to possible mechanisms underlying the brain structural changes, and then tested the emerging putative mechanisms in an experimental model.

**Figure 1.**
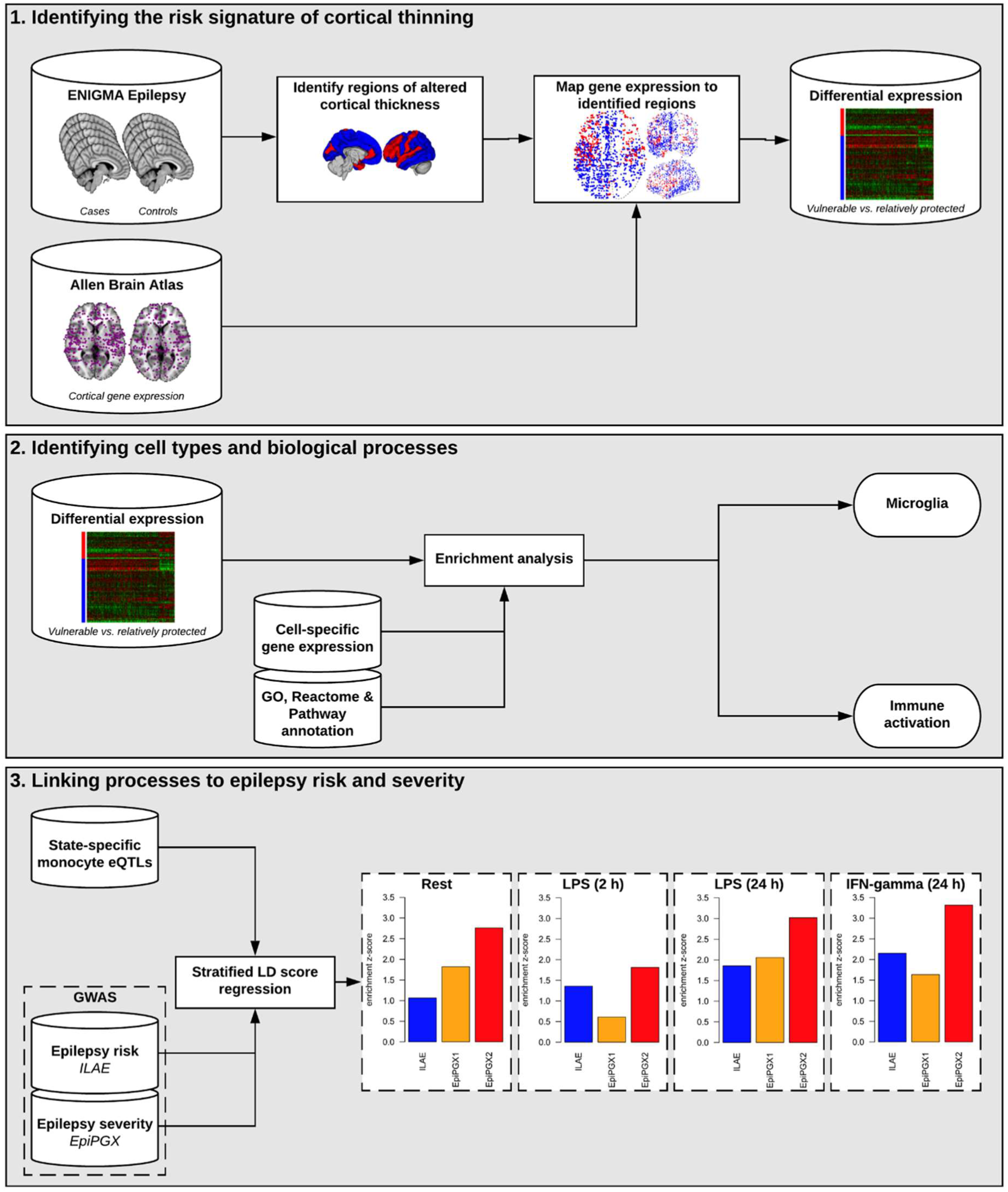
Analysis overview. **Panel a:** Cortical samples of the AIBS dataset were marked as “vulnerable” or “relatively protected” based on whether they were located in a brain region that showed significant reduction in cortical thickness or not, respectively, in a large imaging study^8^. Differential gene expression analysis between the two groups produced the genetic risk signature of reduced cortical thickness in epilepsy. **Panel b:** Significantly differently expressed genes with higher expression in “vulnerable” regions were found to be enriched for marker genes for microglia as well as immune activation related pathways. **Panel c:** LD score regression estimating the enrichment of immune response eQTL signatures in different epilepsy GWAS finds strong enrichment in disease severity (drug-resistant vs drug-susceptible) but not in disease risk (cases vs controls).

The AIBS healthy human brain gene expression atlas provided gene expression profiles for all cortical regions of interest. This gene expression atlas is unique in that from 158 to 348 different brain structures have been sampled from each individual (n=6) covering in total 414 different structures, and that sampling has been precisely mapped to MNI space, enabling linkage of the expression data to external brain maps in MNI space. Thus, the gene expression profiles used in this analysis were highly regionally specified. Of the 34 cortical regions profiled within the ENIGMA-Epilepsy data set, 11 were considered vulnerable to reduced cortical thickness while 23 were considered relatively protected^8^. We compared these two types of cortical regions for significant differences in gene expression and identified 3,258 genes at the chosen P-value threshold (Supplementary Table 1). Interestingly, this gene list was dominated by the over-expression of genes within vulnerable regions (77.3%), with only 22.7% of the genes identified being more highly expressed in relatively protected cortical regions. Eleven genes (*ATXN7L1, C12orf5, LDLRAD4, LRPAP1, NTM, RBBP4, SH3PXD2A, SLC44A5, SPECC1, UBE3C, ZNF81*) had isoforms that were both over- and underexpressed. Considered together, these findings suggest that vulnerability to epilepsy-related reduced cortical thickness is primarily driven by specific regional risk factors, rather than the relative absence of protective factors.

### Genes associated with reduced cortical thickness in the common epilepsies are significantly enriched for microglial markers

Although all cortical regions are similar in structure and cellular composition, there are regional differences, which are detectable within bulk gene expression data^20–23^. We determined whether the genes differentially expressed in vulnerable, as compared to relatively protected, cortical regions could be used to infer cell type(s) involved in mediating cortical thinning (Figure 1). First, we made use of cell-specific gene expression data generated using bulk mouse cortex, and covering the major cellular classes, namely astrocytes, endothelial cells, microglia, myelinating oligodendrocytes, newly-formed oligodendrocytes, oligodendrocyte precursor cells and neurons^24^. We used murine data initially, because of its more robust nature, unconfounded by disease effects. Using this approach, we identified a significant enrichment of sets of microglial-marking genes amongst those genes overexpressed in vulnerable cortex (P_FDR_=1.01×10^-26^; Supplementary Table 2). Conversely, those genes overexpressed in relatively protected cortical regions were enriched for sets of neuron-specific genes (P_FDR_=2.34×10^-6^; Supplementary Table 3). The availability of comprehensive, mouse brain single-cell RNA sequencing data^25^ allowed us to investigate these findings further. Using these independently-derived data, we confirmed the significant enrichment of sets of microglial-marking genes, as well as genes enriched within perivascular macrophages and myocilin-expressing astrocytes, amongst those overexpressed in vulnerable cortex (Figure 2, Supplementary Table 4). In contrast, genes overexpressed in relatively protected cortical regions were strikingly enriched for gene sets specific to *Vglut1*-positive excitatory neurons (Figure 2, Supplementary Table 4).

**Figure 2.**
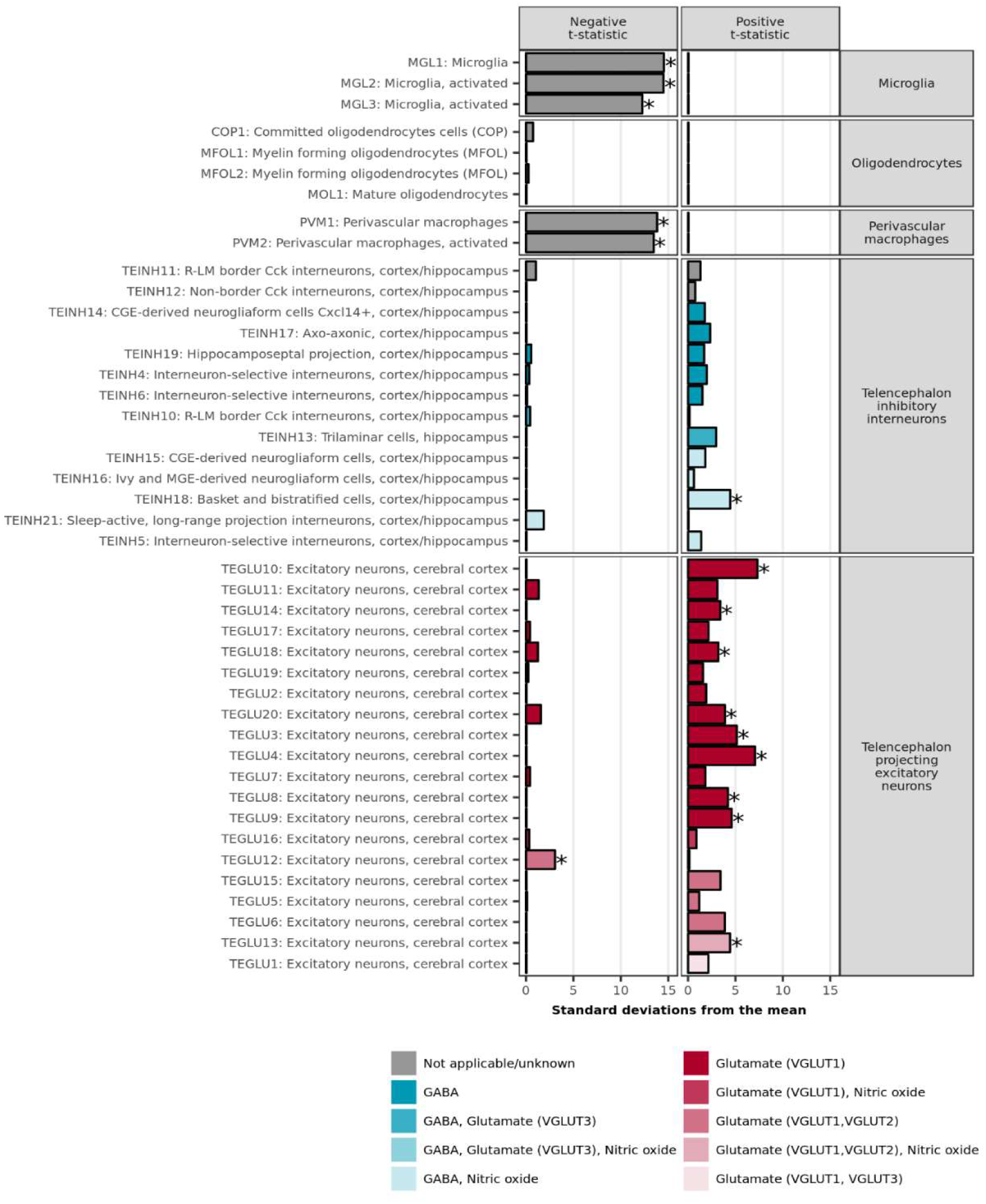
Cell type-specific enrichments amongst genes overexpressed in “vulnerable” and “relatively protected” cortical regions. Using EWCE, genes passing an FDR<0.01 with a negative or positive t-statistic, (namely genes more highly expressed in “vulnerable” as opposed to “relatively protected” cortical regions respectively) were tested for their enrichment in various CNS-associated neuronal and glial clusters identified by Zeisel et al^25^ in a murine single-cell dataset. Displayed here are all clusters whose origin is within cortical regions (amounting to 43 of the 127 clusters tested in total). Neuronal clusters are coloured by their associated neurotransmitter, as determined by Zeisel et al. using expression of genes coding for neurotransmitter transporters or enzymes crucial to their synthesis. Standard deviations from the mean indicate the distance of the mean expression of the target list from the mean expression of the bootstrap replicates. Asterisks denote significance at P_FDR_<0.05 after correcting for multiple testing with the FDR method over all gene sets and cell types displayed. For a full list of all clusters, gene sets, and numerical results see Supplementary Table 4.

Since evidence suggests species differences in cell-specific expression^26^, we extended these findings using human-derived data^26^. Although these human data are more prone to confounding errors due to the small number of tissue samples used and the fact that cortex samples were obtained from surgical resections for a range of brain diseases including epilepsy (18 of 22), we were able to confidently replicate our major findings. In relatively protected areas, we confirmed enrichment of a neuron marker gene set derived from human tissue (P_FDR_=2.77×10^-43^; Supplementary Table 3). We also identified a highly significant enrichment of microglia-marking genes amongst those overexpressed in vulnerable cortical regions (P_FDR_=6.82×10^-54^; Supplementary Table 2). Together, these analyses from mouse and human data suggest that vulnerability to reduced cortical thickness is likely to be mediated at least partly through microglia-specific processes.

### Evidence for selected processes and microglial signatures amongst genes associated with reduced cortical thickness in the common epilepsies

Moving beyond individual genes and cell types, we next investigated the biological processes that could underpin regional differences in vulnerability to reduced cortical thickness in epilepsy. In the first instance we investigated differentially expressed genes for evidence of Gene Ontology, KEGG and REACTOME pathway enrichment. Amongst genes overexpressed in relatively protected cortical regions, the most significant terms we identified related to synaptic function (GO: Synapse, P_FDR_=1.60×10^-7^; GO: Neuronal postsynaptic density, P_FDR_=7.27×10^-5^; Supplementary Table 5). Conversely, amongst genes overexpressed in vulnerable cortical regions, the most significant terms related to immune function and regulation (GO: Immune system process, P_FDR_=6.68×10^-14^; GO: Positive regulation of immune system process, P_FDR_=2.16×10^-13^; Supplementary Table 6), consistent with our results derived using cell-specific gene sets. However, this approach also enabled us to obtain more specific process-related information and suggested the importance of the interferon gamma signalling pathway (GO: Cellular response to interferon gamma, P_FDR_=4.80×10^-8^; GO: Response to interferon gamma, P_FDR_=9.08×10^-8^; Supplementary Table 6).

Since microglia can exist in a range of activation states within the context of epilepsy^27^, we sought to integrate the cell-specific and pathway analyses already performed, to identify the microglial cell states of greatest importance in reduced cortical thickness. We collated gene signatures for distinct microglial states from the existing literature and also inferred signatures of microglial state through co-expression network analyses (see Online Methods). While there were significant overlaps in gene membership across the 16 microglial signatures used (Supplementary Figure 1), each of the gene lists were distinct. We identified a significant enrichment for 10 of the 16 signatures with genes overexpressed in vulnerable cortex. This included the microglial signature generated by Srivastava and colleagues^28^, which was positively correlated with seizure frequency in a mouse model of chronic epilepsy (P_FDR_=4.81×10^-17^; Supplementary Table 6). Striking enrichments were also identified for an inferred human microglial signature enriched for type 1-like microglial markers (P_FDR_=2.53×10^-42^; grey60; Supplementary Table 6), as well as signatures for aged, late activation, and de-activated microglia. However, we saw little evidence for enrichment within signatures of early activation (“Early Response” signature^29^, P_FDR_=4.53×10^-1^). Given that these enrichment results are ultimately derived from humans with an average duration of epilepsy of 17.42 (+/-11.99) years, the results suggest that the microglial states contributing to reduced cortical thickness in epilepsy of this duration are shared with those microglial states recently implicated in various forms of chronic neurodegeneration^29–31^. However, the data do not exclude contributions from other microglial states at early stages of epilepsy in humans, including during epileptogenesis.

### Genetic evidence in support of immune activation as a modifying, but not causal, factor in common epilepsies

Region-specific reduced cortical thickness across the common epilepsies was correlated with disease duration and age of onset of epilepsy^8^. Given that the analyses described above suggest the importance of microglia responses in vulnerability to reduced cortical thickness, we predicted that genetic variants affecting microglial responses would also impact upon the severity of epilepsy. While no microglial eQTL data set exists to date, eQTL analyses have been performed using monocytes at rest, and also monocytes treated with IFN-γ or lipopolysaccharide (LPS)^32^. We postulated that these eQTLs would be enriched for heritability of risk loci for epilepsy that is drug-resistant (one surrogate for disease severity) but not for epilepsy *per se*. To investigate this, we used GWAS data on epilepsy susceptibility^2^ and separate GWAS data on the phenotype of drug-resistant epilepsy from the EpiPGX consortium (www.epipgx.eu). Using LD score regression, we sought enrichment in heritability for these phenotypes within state-specific monocyte eQTLs^32^. We found no significant enrichment in the heritability of epilepsy in this form of annotation, suggesting that susceptibility to common epilepsies is less likely to be driven by immune processes (Table 1). However, eQTLs regulating the response to IFN-γ were highly enriched for drug-resistant epilepsy vs drug-responsive epilepsy loci (P=0.00045; P_FDR_=0.0095, Table 1). This result was particularly striking given the small size of this GWAS (2423 drug-resistant cases vs 1626 drug-responsive cases). Thus, we provide genetic evidence in support of microglial-mediated responses as a modifying factor in severity of epilepsy, but not its susceptibility.

### Widespread regionalised over-representation of microglia in human brain tissue from people with epilepsy

Based on the analyses above, we predicted that brain tissue from individuals with various forms of epilepsy would have regionalized higher densities of microglia as compared to tissue from non-epilepsy controls and that this would be apparent beyond the context of acute seizure activity. In order to test this hypothesis, we investigated microglia density across 14 regions of interest (ROIs) in *post mortem* brain tissue derived from 55 individuals, including individuals with non-lesional epilepsy (EP-NL, n=18), lesional epilepsy (EP-L, n=21) and non-epilepsy controls (NEC, n=16). Using Iba-1 immunolabelling, we found that single ramified microglia and processes were found throughout the cortex; scattered perivascular macrophages were also labelled (Figure 3a). Enlarged and more complex/branching microglia, focal aggregates and amoeboid/macrophage forms were noted in some ROIs (Figure 3a). Consistent with our hypothesis, the Iba1 labelling index was significantly higher in all epilepsy (EP-NL and EP-L together) than NEC for all ROIs (P=4.0×10^-13^) and for both subgroup comparisons (EP-NL [P=3.7×10^-13^] and EP-L [P=3.5×10^13^]) against NEC. Regional differences were noted within the epilepsy groups, and for individual ROIs compared between epilepsy and control groups (see Figure 3 and Supplementary Table 7). Whilst we did not attempt to formally map histopathological ROIs to the ENIGMA-Epilepsy ROIs because of tissue processing-related artefacts, we noted that the Iba1 labelling index was similar across EP-NL, EP-L and NEC in BA17, as compared to pulvinar and BA22 where the Iba1 labelling index was higher in EP-NL and EP-L groups than in NEC. Thus, these results are consistent with the view that there is an over-representation of microglia in brain tissue from people with chronic epilepsy, and that such microglial responses may occur in a regionally-specific manner.

**Figure 3.**
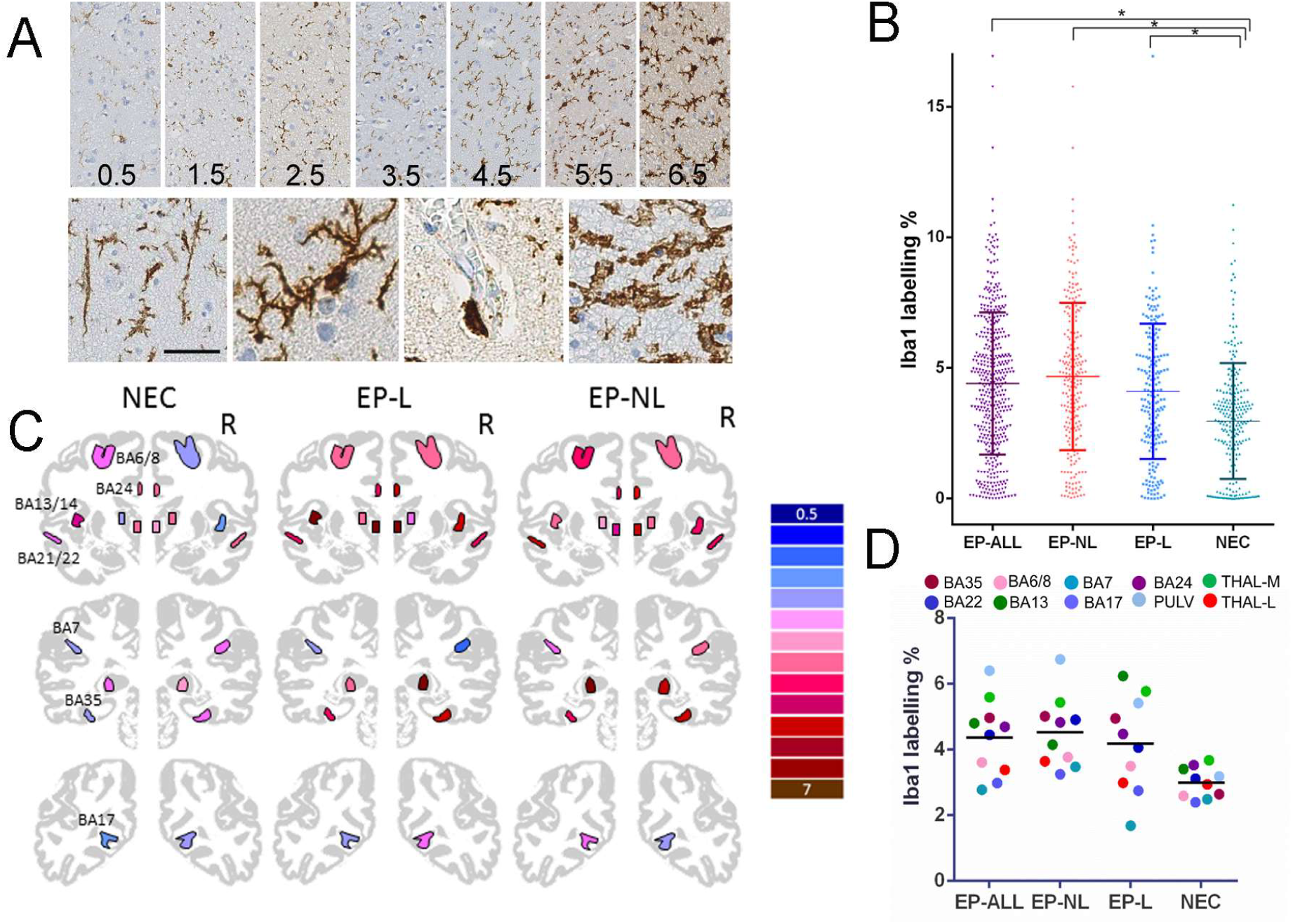
Presence of excess activated microglia in post mortem brain tissue from people with epilepsy. **Panel a:** Top row: Iba1 labelling randomly selected from cases in the study (representing all three groups) with a range of staining from 0.5 to 6.5% with progressive increase in complexity, number and size of ramified microglial (all taken at ×20). Bottom row: High magnification of morphological types of Iba1-labelled cells including (left to right) ‘rod’ cells, ramified microglia, perivascular macrophage and amoeboid forms. Fixation time in illustration of ramified microglia was 467 days (bar = 30 microns). **Panel b:** Scatter graph of all data points from 709 sections including all brain regions showing mean and standard deviation in the four main groups: Epilepsy-ALL (EP-ALL), Epilepsy Non-Lesional (EP-NL), Epilepsy-Lesional (EP-L) and non-epilepsy controls (NEC). EP-ALL, EP-NL, EP-L are all significantly greater than NEC (*respective P-values: 4.0×10^-13^, 3.7×10^-13^, 3.5×10^-13^) **Panel c.** Iba1 immunolabelling shown in 10 Brodmann areas in each hemisphere colour coded for the mean percentage labelling in each group. THAL-M, medial thalamus, THAL-L, lateral thalamus, PULV, lateral pulvinar. **Panel d.** Scatter graph of the mean Iba1 immunolabelling in the same Brodmann areas including both hemispheres in the four groups listed above.

### Experimental evidence in support of microglial activation as a modifying factor for cortical thickness in a mouse model of epilepsy

To provide proof-of-concept evidence causally linking microglia activation to cortical thinning, we used a mouse model of acquired epilepsy where convulsive seizures originate and spread in the limbic system, and also involve the neocortex.^33–36^ This model mimics features of mesial temporal lobe epilepsy with hippocampal sclerosis (MTLE), with neuronal damage also observed in extrahippocampal areas^33,34,36^, as also shown in human MTLE in the ENIGMA-Epilepsy study^8^. Spontaneous seizures are triggered by status epilepticus (SE) evoked in the amygdala. Spontaneous seizures develop after a latent phase after the primary insult (onset, 6.2±0.5 days, n=21) and recur for months (Supplementary Figure 2), and are drug-resistant^34^. Microglia are morphologically activated and proliferate by 3-fold on average within one week after SE as assessed in a different cohort of mice by analysis of Iba-1-positive cells in the forebrain (number of cells, hippocampus: sham, 1013±129; 1 week post-SE, 2837±250, P=0.0001, n=10-11 mice).

First, we studied whether the thickness and volume of selected cortical brain regions were reduced, and those of the lateral ventricles increased (directions of change as predicted by the ENIGMA-Epilepsy findings in humans) in placebo-diet fed epileptic mice (n=10) *vs* sham mice (not exposed to SE, n=8) as assessed by post mortem MRI. We found that the lateral ventricles were enlarged by 2-fold (P=0.0004, one-tailed t-test): this effect was associated with an average 10% significant reduction in the volume of the hippocampus (P=0.0043) and entorhinal cortex (P=0.0061), and caudato-putamen (P=0.049) (Supplementary Figure 3). No significant changes were observed in other brain areas such as the thalamus and the perirhinal cortex, although the average volumes of both were lower than the corresponding values in sham mice (Supplementary Figure 3). Notably, a significant reduction in the thickness of both entorhinal (P=0.0001; Figure 4a) and perirhinal cortices (P=0.0338; Supplementary Figure 4) was also measured in the same mice.

**Figure 4.**
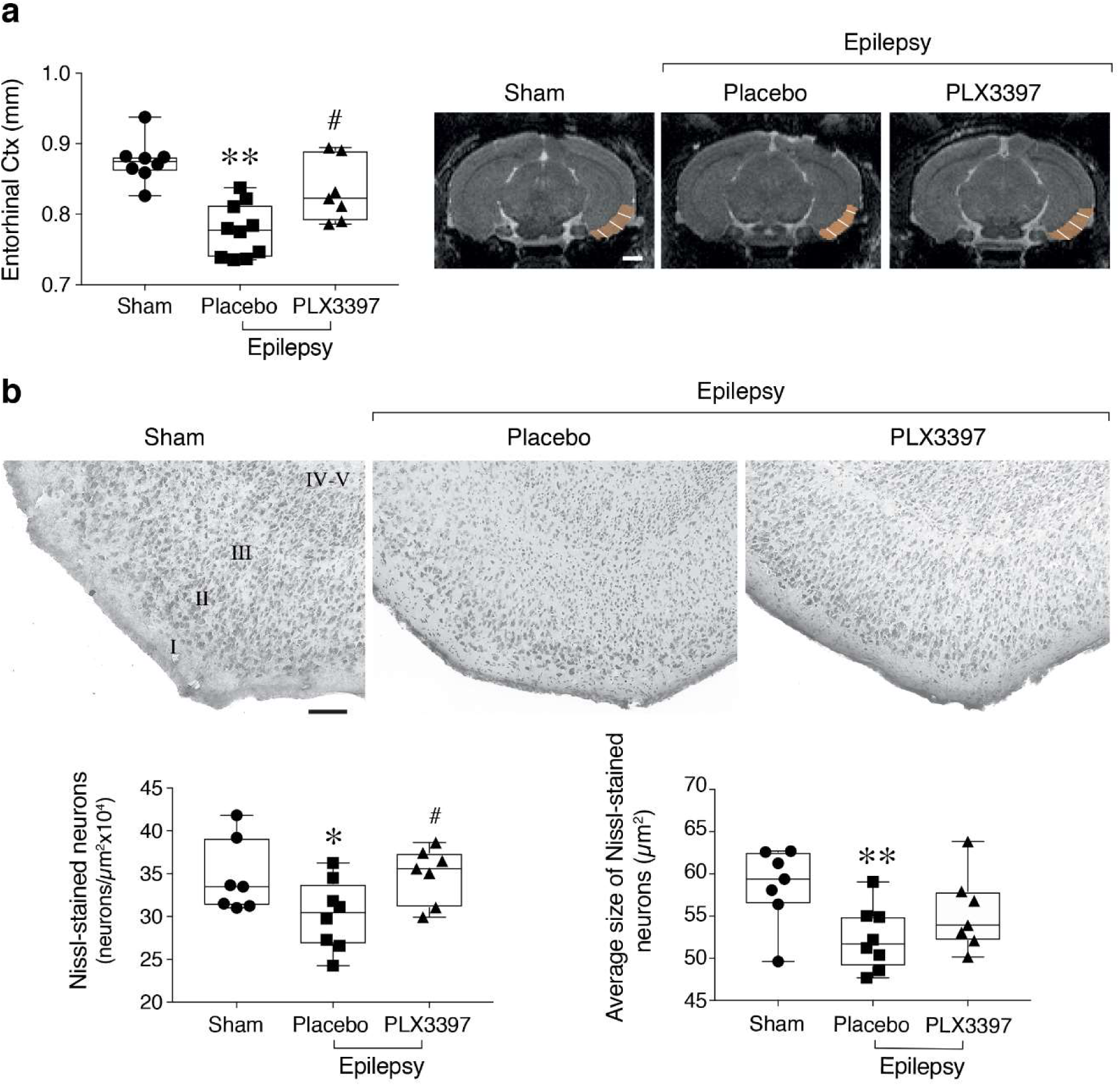
Effects of PLX3397 on entorhinal cortex thickness in epileptic mice. **Panel a:** box-and-whisker plots depicting median, minimum, maximum and single values related to the quantification of entorhinal cortex thickness, as assessed by quantitative postmortem MRI performed at the end of EEG monitoring (placebo: n=10; PLX3397: n=7), and in sham mice (n=8). Mice studied with MRI were randomly selected from those described in Supplementary Figure 2. MRI images depict representative slices showing the ROI used to quantify the cortical thickness. The white line within the ROI was manually drawn to measure the cortical thickness. **P=0.0001 vs sham; #P=0.0068 vs placebo by one-tailed t test. Scale bar: 1 cm. **Panel b:** representative Nissl-stained sections of the entorhinal cortex in the experimental groups (top row), and the relative quantification of the number and the average size of Nissl-stained neurons (bottom row). One sham mouse and two placebo mice were excluded from the analysis due to poor quality of Nissl staining. Data are shown by box-and-whisker plots depicting median, minimum, maximum and single values *P=0.0487, **P=0.0103 vs sham; #P=0.02 vs placebo by one-tailed t test. Scale bars: 100 μm.

Next, we measured brain region volumes and cortical thickness in mice treated with PLX3397. The drug was administered via food to massively reduce the number of microglial cells in the forebrain during the post-insult epileptogenesis phase (prodromal to epilepsy onset) and in the early phase after spontaneous seizures begin. Importantly, this decrease was induced transiently in order to assess the impact of microglia activation in the initial phases of the disease on the structural brain changes we detected in the epileptic mice. We found that the thickness of the entorhinal cortex of microglia-depleted mice was similar to sham mice, thus indicating that microglia depletion prevented the cortical thinning of this ictogenic brain area in epileptic mice (P=0.0068 vs placebo; Figure 4a). Microglia depletion did not affect the thinning of the perirhinal cortex (Supplementary Figure 4) or any of the brain volume changes occurring in epileptic mice (Supplementary Figure 3). Evaluation of neuronal cell numbers in the entorhinal cortex showed a 14% decrease in cell density (P=0.0487 vs sham; Figure 4b) and in their average cell body size (P=0.01 vs sham; Figure 4b) in the epileptic mice receiving placebo diet. The reduction in cell density was prevented in epileptic mice when microglia had been depleted during epileptogenesis by PLX3397 (P=0.02 vs placebo; Figure 4b).

## Discussion

The epilepsies are complex conditions with multiple facets including various causes, differing responses to treatment, and unpredictable outcomes. Most attention has been paid to causation and processes of epileptogenesis across the broad constituent spectrum of syndromes. In contrast, disease progression has not been a primary focus of research even though for some rare epilepsies (the developmental and epileptic encephalopathies, DEE), the window of opportunity to ameliorate disease may be open for longer than expected, as disrupted patterns of gene expression last into adulthood^18^. Here, we show that across the broad spectrum of the more common epilepsies (specifically excluding DEE), patterns of gene expression spatially correlated with reduced cortical thickness implicate neuroinflammatory processes, and, particularly, microglial activation, as contributors to the underlying cause of reduction in cortical thickness. We show that this molecular signature of neuroinflammation significantly overlaps with a gene set already causally linked to seizure frequency in a mouse model of chronic epilepsy^28^. Further, our data add new evidence showing that microglial activation is associated with at least some of the structural changes seen in brain areas involved in seizure circuitry, in that microglial depletion can directly prevent associated cortical thinning.

We tested the hypothesis that microglial/monocyte activation is a key modulator of the severity of epilepsy using both genetic and functional approaches. We used the availability of GWAS data for resistance to anti-epileptic drug treatment, a marker of disease severity, to investigate enrichment in heritability at genetic loci already known to influence the expression of genes involved in monocyte activation. We note this analysis assumes a high degree of similarity in the genetics of gene expression in monocytes and microglia. We find a highly significant enrichment in the heritability of epilepsy severity amongst these immunomodulatory loci, despite the absence of a significant enrichment for heritability of epilepsy per se. Finally, in keeping with these observations, experimental microglial depletion timed to coincide with a period of epileptogenesis in a murine model can prevent regionalised cortical thinning, but does not influence the eventual development of seizures themselves in our model. Data from the experimental model also demonstrate that the cortical thinning is at least partly due to reduced neuronal cell density and average neuronal size: reduction of neuronal density can be prevented by appropriately-timed microglial depletion (neuronal size changes were not rescued, which may explain the observation that entorhinal volume changes are not completely prevented by microglial depletion). We, and others^37–41^, find activated, Iba1^+^-microglia are present in excess in brain tissue from people with epilepsy, compared to brain tissue from healthy controls, providing evidence for translation to human epilepsies of our assertions from the experimental model data. Together, these findings separate important processes occurring in the course of the epilepsies, and incriminate potentially modifiable microglial activation states in the hitherto largely-ignored feature of cortical thinning in the common human epilepsies. Our results also suggest other factors are likely to be involved.

Importantly, we note that the clue to the possible role for microglial activation in cortical thinning came from a cross-sectional study of chronic human epilepsy: although reduced cortical thickness correlated with disease duration^8^, we could not distinguish whether the structural difference had developed at disease onset (e.g. with causation), during epileptogenesis, during the course of the disease, or a combination of these epochs. Our multimodal data, and especially the experimental model results, allow us to begin to address this question. The model relates to early processes in epileptogenesis, and show a separation for microglial roles in cortical thinning and seizure occurrence, whilst data from another model relates to the chronic disease state, and shows an effect of microglial manipulation on seizure frequency in that chronic state^28^ (cortical thinning was not assessed in that model). The models and data are not directly compatible, and we cannot test hypotheses arising from the chronic model in data from ENIGMA-Epilepsy. However, taken together, the data suggest microglia may have multiple modifying roles during epileptogenesis and progression of disease across common human epilepsies, though we find no evidence that they contribute to the actual occurrence of these common forms of epilepsy (from either our human or murine data). That seizure frequency and cortical thinning may be separable processes adds to important epidemiological evidence that seizure frequency is not the only contributor to morbidity in people with a history of epilepsy.^5^ Microglia are already known to have many roles in specific types of epilepsy, demonstrated clearly in a variety of animal models. Such roles include phagocytosis, which may link consumption of synapses with cognitive changes in long-term active epilepsy,^42^ providing another possible mechanism for actual loss of brain volume in epilepsy: ‘time is brain’^43^.

Considering the complex relationship between inflammation and epilepsy, it is noteworthy that despite the growing evidence for neuroinflammation as a modulator of epilepsy, overt neuroinflammation is considered causal in a very few, rare, severe human epilepsies. Rasmussen’s encephalitis is a severe drug-resistant epilepsy characterised by progressive loss of cerebral substrate, in which astrogliosis and T-lymphocyte infiltration are central features, with IFN-γ and granzyme B overactivity and evidence of efficacy of α4 integrin blockade^44^. Microglial activation is also seen in Rasmussen’s encephalitis, and in tissue from epilepsies due to hippocampal sclerosis and mesial temporal lobe sclerosis, focal cortical dysplasia (FCD) and tuberous sclerosis^37–41^. The latter two conditions are known to have genetic, rather than inflammatory, causes, but the extent of microglial activation in the chronic disease phase correlates with seizure frequency and disease duration in these studies^37,41^, pointing again to distinction between processes related to the initial cause and others manifest during active disease. Importantly, resected human brain tissue is only available from a few cases of a few types of epilepsies (mostly surgical specimens from MTLE and FCD^45^), so that it is impossible to otherwise evaluate the role of microglia using neuropathological data in the majority of common human epilepsies, from which brain tissue cannot be obtained in life. Brain imaging in animal models using a label (TSPO) for microglial activation shows dynamic upregulation during epileptogenesis, with persistent, although declining, activity in the chronic phase and correlation with spontaneous seizures^46^; in chronic human TLE, there is increased TSPO binding ipsilateral and contralateral to seizure foci^47^. Using immunolabelling, we demonstrate that there is over-representation of activated microglia in human and experimental brain tissue, compared to controls. However, neither TSPO imaging nor neuropathological study is realistically applicable to large numbers of people with epilepsy, whilst MRI is. We propose, using MRI-derived patterns and correlation with gene expression, that microglial-associated neuroinflammation drives the important, but as yet largely neglected, phenotype of cortical thinning in a broad swathe of common human epilepsies. Subsequent experimental intervention in an animal model suggests that early manipulation of microglia has the capacity to rescue disease-related cortical thinning, opening up new areas for attention and treatment in common human epilepsies.

Our results point to important roles for neuroinflammation and potentially specific molecular actors, such as IFN-γ. However, the diversity of microglial states and functions, and the complex, dynamic, interactions between neurons, astroglia and microglia, that at the very least can promote epileptogenesis^27^ have yet to be fully resolved. Our key observation is that microglial activation is not a rare phenomenon in the epilepsies, but is likely to be central across the breadth of types of epilepsy. Not only does this deepen understanding and show that some disease processes and progression in epilepsy may be independent of cause, but also offers new avenues for treatment and disease modification. Manipulation of microglial activity may not only address the common phenomenon of treatment resistance^28^, but may also prevent irreversible loss of brain substance.

